# The origin of animal multicellularity and cell differentiation

**DOI:** 10.1101/161695

**Authors:** Thibaut Brunet, Nicole King

**Affiliations:** Howard Hughes Medical Institute and the Department of Molecular and Cell Biology, University of California, Berkeley, CA

**Keywords:** multicellularity, choanoflagellates, Choanozoa, metazoan origins

## Abstract

How animals evolved from their single-celled ancestors over 600 million years ago is poorly understood. Comparisons of genomes from animals and their closest relatives – choanoflagellates, filastereans and ichthyosporeans – have recently revealed the genomic landscape of animal origins. However, the cell and developmental biology of the first animals have been less well examined. Using principles from evolutionary cell biology, we reason that the last common ancestor of animals and choanoflagellates (the ‘Urchoanozoan’) used a collar complex - a flagellum surrounded by a microvillar collar – to capture bacterial prey. The origin of animal multicellularity likely occurred through the modification of pre-existing mechanisms for extracellular matrix synthesis and regulation of cytokinesis. The progenitors of animals likely developed clonally through serial division of flagellated cells, giving rise to sheets of cells that folded into spheres by a morphogenetic process comparable to that seen in modern choanoflagellate rosettes and calcareous sponge embryos. Finally, we infer that cell differentiation evolved in the animal stem-lineage by a combination of three mechanisms: division of labor from ancient plurifunctional cell types, conversion of temporally segregated phenotypes into spatially segregated cell types, and functional innovation.

## Introduction

Every aspect of animal life – from morphology to physiology and behavior – requires the cooperation of thousands to billions of cells. This multicellular state is established in each generation by serial divisions of a single cell, the zygote. Under joint control by the genome and the environment, daughter cells produced by these divisions change shape, migrate, and selectively attach or detach to give rise to the adult body form through a process known as morphogenesis. In parallel, a process of cell differentiation under fine spatiotemporal control delineates the division of labor between each of the final cell types. The correct execution of this cellular choreography, repeated anew in every generation, is fundamental to the life of every animal on the planet.

Yet, this type of complex development did not always exist. The discontinuous phylogenetic distribution of multicellularity and fundamental differences in cellular mechanisms argue that multicellularity evolved independently in at least 16 different eukaryotic lineages, including in animals, plants, and fungi (Gould 1996; Bonner 1998; King 2004; Rokas 2008; Knoll 2011). Thus, the mechanisms underpinning animal multicellularity and spatially controlled cell differentiation were likely elaborated in the stem lineage of animals, building upon pathways present in their single-celled ancestors (Richter and King 2013).

Despite the centrality of multicellularity and cell differentiation to animal biology, their origins are relatively poorly understood. What did the single-celled ancestors of animals look like? How and when did multicellularity and cell differentiation evolve, and in what ecological context? What were the underlying molecular mechanisms? Did features of the single-celled progenitors of animals facilitate the early evolution of multicellularity? Conversely, did this single-celled ancestry exert constraints upon the form and function assumed by early animal ancestors?

While the gene complements of animal ancestors have been discussed in depth elsewhere and will not be the focus of this review (see e.g. (King 2004; King et al. 2008; Larroux et al. 2008; Richter and King 2013; Suga et al. 2013; de Mendoza et al. 2013; Sebé-Pedrós et al. 2017)), it is notable that many genes required for animal multicellularity (e.g. tyrosine kinases (King and Carroll 2001; King et al. 2003), cadherins (Abedin and King 2008; Nichols et al. 2012), integrins (Sebé-Pedrós et al. 2010; Suga et al. 2013), and animal-like extracellular matrix domains (King et al. 2008; Williams et al. 2014)) evolved before animal origins. Building upon these ancient proteins, the stem-animal lineage was notably marked by an explosive diversification of transcription factor families and signaling molecules, fueled both by the emergence of new families (e.g. the Antennapedia and Pax families of homeodomain transcription factors and the secreted signaling proteins Wnt and BMP) and by expansion of existing families (Larroux et al. 2008; Srivastava et al. 2010; de Mendoza et al. 2013). The cell biology and morphology implemented by these ancestral genomes, however, have been less thoroughly explored (Nielsen 2008; Richter and King 2013; Arendt et al. 2015; Cavalier-Smith 2017). In this review, we will consider how the evolution of cellular phenotype shaped the origin of animals.

Although the first animals evolved over 600 million years ago, meaningful insights into their origin may be gained through the comparison of extant lineages. This approach has revealed a number of features that were likely present in the last common ancestor of animals, the “Urmetazoan” (Figure 1). For example, nearly all extant animals have obligate multicellularity (see (Metzger et al. 2015; Chang et al. 2015) for exceptions in some parasitic forms) with adult stages typically displaying a specialized morphology and at least five morphologically distinguishable cell types (Valentine 2006; Smith et al. 2014). This suggests that the Urmetazoan evolved from a lineage with a long prior history of obligate multicellularity (Nielsen 2008; Richter and King 2013; Budd and Jensen 2017). Likewise, multicellularity in animals is invariably the result of a complex embryogenesis initiated by sperm/egg fusion, followed by serial rounds of cell divisions (cleavage). This suggests the same was true of the first animals. Finally, in every major animal lineage from sponges and ctenophores to bilaterians, the cells of the future feeding cavity move inside the embryo, morphogenesis establishes the adult body shape, and cell differentiation takes place (Leys and Degnan 2002; Arendt 2004; Leys and Eerkes-Medrano 2005; Ereskovsky 2010; Nielsen 2012). A form of this elaborate developmental process presumably already existed in the Urmetazoan. Rather than evolving in one step in a single-celled ancestor, it more plausibly resulted from a long and gradual evolution. Therefore, to more fully reconstruct the origin of animal development, we must extend our comparisons beyond animals to include their closest living relatives.

**Figure 1.**
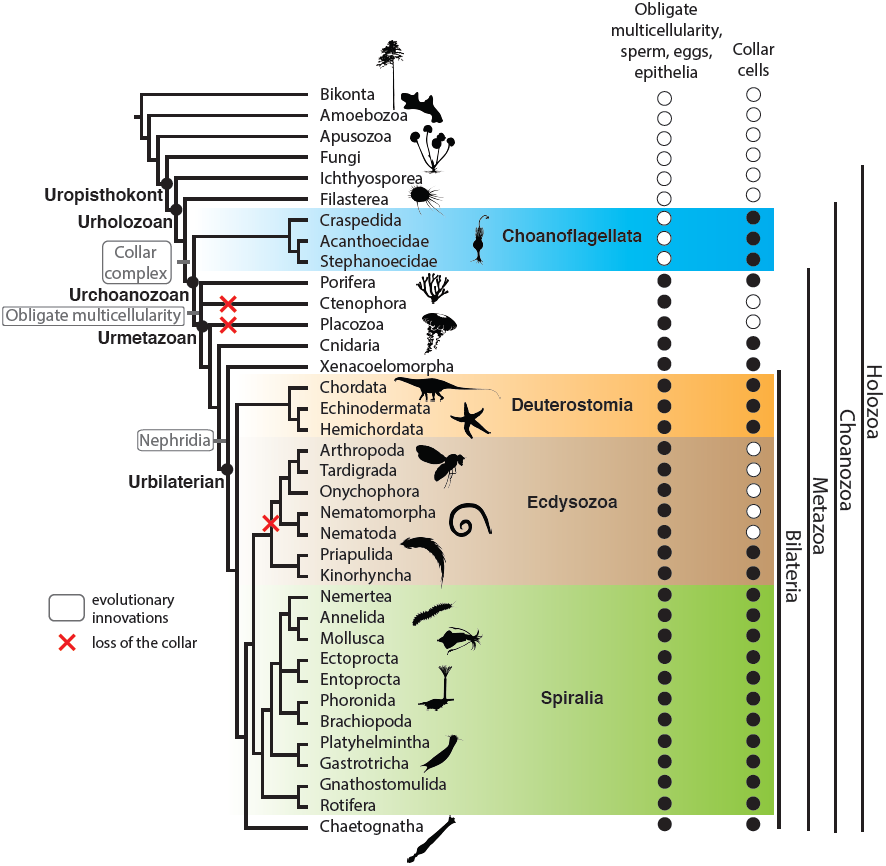
Phylogenetic distribution of traits inferred in the Urmetazoan. The presence of epithelia (Leys et al. 2009), sperm, eggs and multicellularity (Nielsen 2012) and collar complex (see Supplementary Table 1 for details and references) are mapped onto a eukaryotic phylogeny. The collar complex is inferred to have been present in the Urchoanozoan, and to be a choanozoan synapomorphy. The eukaryotic phylogenetic tree follows (Torruella et al. 2015). The animal phylogenetic tree is a consensus based on (Struck et al. 2014; Laumer et al. 2015) for Spiralia, (Borner et al. 2014) for Ecdysozoa, (Cannon et al. 2016) for Xenacoelomorpha and (Telford et al. 2015) for other animal phyla. Species silhouettes are from PhyloPic (http://phylopic.org).

The sister-group of animals has unambiguously been shown to be the choanoflagellates (King et al. 2008; Ruiz-Trillo et al. 2008; Shalchian-Tabrizi et al. 2008; Torruella et al. 2015), a globally distributed group of marine and freshwater protists (Leadbeater 2014) with a highly distinctive morphology (Figure 2A,B,E). Choanoflagellates are characterized by an apical flagellum surrounded by a collar of microvilli, which together form a “collar complex” (Leadbeater 2014). The beating of the flagellum generates forces that can both propel the cell through the water column and produce a flow that allows the choanoflagellate to collect bacterial prey on the outer surface of the microvillar collar. The morphological similarity between choanoflagellates and certain animal cells, particularly sponge choanocytes, was evident even to the first choanoflagellate observers (James-Clark 1867) and inspired the hypothesis of a close relationship between choanoflagellates and animals. This hypothesis, first proposed in the 19^th^ century on morphological grounds, remained otherwise untested for more than a century, as both the affinities of choanoflagellates to sponges and of sponges to other animals were the subject of a variety of alternative hypotheses (see historical summary in (Leadbeater 2014)). The issue was only settled with the application of comparative genomics and molecular phylogenetics, which provided conclusive evidence both that sponges belong to the animal kingdom (Wainright et al. 1993; Srivastava et al. 2010; Telford et al. 2015) and that animals and choanoflagellates are sister-groups (King et al. 2008; Ruiz-Trillo et al. 2008; Shalchian-Tabrizi et al. 2008; Torruella et al. 2015). Together, choanoflagellates and animals thus form a monophyletic clade, which we refer to as “Choanozoa” (see box). Furthermore, phylogenomic studies of previously enigmatic taxa have revealed the closest living relatives of choanozoans (Ruiz-Trillo et al. 2008; Torruella et al. 2015) to be Filasterea (a group comprising filopodiated amoebae and recently discovered flagellated protozoans (Hehenberger et al. 2017)) and Ichthyosporea (which alternate between large syncytial spores and individual amoebae). Together, Choanozoa, Ichthyosporea, and Filasterea make up the clade Holozoa, which is the sister-group of Fungi (Figure 1).

**Table 1.** Cell cycle and ECM genes from diverse eukaryotes regulate the unicellularity/multicellularity switch.

| Species | Type of multicellularity | Gene required for the unicellular/multicellular switch | Gene function |
| --- | --- | --- | --- |
| *Gonium pectorale* (green alga) | Clonal | *retinoblastoma* | **Cell cycle** (transcription inhibitor) (1) |
| *Saccharomyces cerevisiae* (fungus) | Aggregative (flocculation) | *flo1, flo5, flo8, flo9, flo10, flo11, sta1* | **ECM** (*flo1, flo5, flo9, flo10, flo11*: lectins, *sta1*: endoamylase, *flo8*: transcriptional activator of the formers) (2-6) |
| *Saccharomyces cerevisiae* (fungus) | Clonal (chain and snowflake mutants) | *cts1*<br>*ACE2* | **ECM** (chitinase) (7)<br>**Cell cycle** (transcription factor) (8,9) |
| *Dictyostelium discoideum* (slime mold) | Aggregative | *cbp-26* | **ECM** (lectin) (11,12) |
| *Salpingoeca rosetta* (choanoflagellate) | Clonal | *rosetteless* | **ECM** (lectin) (10) |

References: 1. (Hanschen et al. 2016); 2. (Douglas et al. 2007); 3. (Lo and Dranginis 1996); 4. (Soares 2011); 5. (Stratford 1992); 6. (Fidalgo et al. 2006); 7. (Kuranda and Robbins 1991); 8. (Oud et al. 2013); 9. (Ratcliff et al. 2015); 10. (Shinnick and Lerner 1980); 11. (Ray et al. 1979); 12. (Levin et al. 2014)

**Figure 2.**
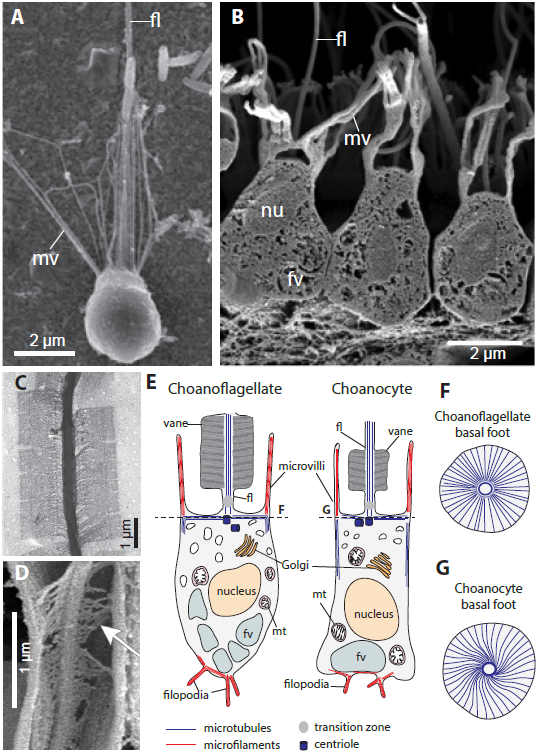
Morphology and ultrastructure of choanoflagellates and sponge choanocytes. (A) Scanning Electron Micrograph (SEM) of the choanoflagellate *Salpingoeca rosetta*, reproduced from (Dayel et al. 2011). In this panel and all others: (fl): flagellum, (mv): microvilli. (B) SEM of choancoytes from the sponge *Sycon coactum*, from (Leys and Hill 2012). In this panel and others: (nu): nucleus, (fv): food vacuole. (C) Transmission Electron Micrograph (TEM) of the flagellar vane of the choanoflagellate *Salpingoeca amphoridium*, from (Leadbeater 2014). (D) SEM of the flagellar vane of a choanocyte from the sponge *Spongilla lacustris*, from (Mah et al. 2014). Arrow shows the fibrous structure of the vane and lateral contact with the collar. (E) Comparative ultrastructural schematics of a choanoflagellate and a sponge choanocyte, modified from (Maldonado 2004) following (Woollacott and Pinto 1995) for the microtubule cytoskeleton and (Karpov and Leadbeater 1998) for the actin cytoskeleton. The presence of filopodia in choanocytes is indicated following (Grassé 1973). (mt): mitochondria. (F,G) basal microtubular foot supporting the flagellum in choanoflagellates and choanocytes, following (Garrone 1969; Woollacott and Pinto 1995; Leadbeater 2014).

### Box 1: Choanozoa: the clade composed of choanoflagellates and animals.

We define Choanozoa as the clade containing the most recent common ancestor of animals (represented by *Homo sapiens* Linnaeus 1758) and choanoflagellates (represented by *Monosiga brevicollis* Ruinen 1938), along with all of its descendants. The Greek root “*choanē*” (or funnel) refers to the collar, which in the current state of knowledge is a synapomorphy of the clade. Although “Choanozoa” was used previously to refer to an assemblage of protists (Cavalier-Smith, 1991) that later proved paraphyletic (Shalchian-Tabrizi et al., 2008); this usage was not adopted and the name is more appropriately applied as defined here. The informal term “choanimal” (Fairclough et al., 2013) and the formal term Apoikozoa (Budd and Jensen, 2017) have both been previously proposed for the clade containing choanoflagellates and animals, but neither has been formally described nor fully adopted. In particular, the term “Apoikozoa” is less fitting, as the root “*apoiko-*” refers to colony formation, which is neither universally present in choanozoans, nor exclusive to them.

## I. Choanoflagellates reveal the cellular foundations of animal origins

In light of the sister-group relationship between choanoflagellates and animals, a fundamental question is whether shared cellular features – such as the collar complex – were already present in the last common choanozoan ancestor, the “Urchoanozoan” (Figure 1). Electron microscopy has revealed that the similarities between choanoflagellates and choanocytes extend beyond morphology to include a shared underlying ultrastructure (Richter and King 2013). In both choanoflagellates and sponge choanocytes, the flagellum is supported by microtubules (Karpov and Leadbeater 1998; Gonobobleva and Maldonado 2009) and often displays a characteristic “vane” – a pair of bilateral wing-like filamentous extensions of the flagellum that have only been reported in choanoflagellates and in sponge choanocytes (Figure 2C,D) (Petersen 1929; Vlk 1938; Hibberd 1975; Mehl and Reiswig 1991; Leadbeater 2006; Mah et al. 2014). The ovoid cell body is encased in parallel arrays of sub-membranous microtubules that emerge from the basal body and span the cell from the apical to the basal side. Underneath the flagellum, the basal body is supported by a basal foot surrounded by an elaborate “crown” of transverse microtubules. Like the flagellar vane, this organization appears unique to sponge choanocytes and to choanoflagellates (Figure 2F,G) (Garrone 1969; Woollacott and Pinto 1995; Leadbeater 2014). Finally, in both choanoflagellates and choanocytes, the microvilli are supported by bundled actin microfilaments maintained at a constant length within a given cell (Karpov and Leadbeater 1998; Rivera et al. 2011). Choanoflagellate genomes encode homologs of most animal microvillar proteins, among which two families appear choanozoan-specific: Ezrin/Radixin/Moesin (ERM) and Whirlin, both of which are involved in controlling microvillar length (Sebé-Pedrós et al. 2013a; Peña et al. 2016). This further supports the notion that the collar is a choanozoan synapomorphy^1^.

Besides its conserved ultrastructure, the stem choanozoan ancestry of the collar complex is further supported by its broad distribution in animals. Beyond sponges, collar complexes featuring a flagellum surrounded by a ring of microvilli are found in most animal phyla (Figure 1, Supplementary Table 1) – e.g. in epidermal cells (often sensory (Supplementary Figure 1A) (Fritzsch et al. 2007; Fritzsch and Straka 2014)), terminal cells of protonephridia ensuring excretion (Supplementary Figure 1B) (Ruppert and Smith 1988), or as part of diverse inner epithelia (Supplementary Figure 1C) (Nerrevang and Wingstrand 1970; Goldberg and Taylor 1989). Collar cells often function in food absorption: choanoflagellates and sponge choanocytes phagocytose live bacteria, and the collar cells lining the gastrodermis of some cnidarians endocytose food particles produced by extracellular digestion (Goldberg and Taylor 1989). In bilaterians and ctenophores, nutrient acquisition through endocytosis is performed by enterocytes lining the midgut that frequently display a motile flagellum and microvilli (although the microvilli are packed into a dense brush border rather than forming a simple ring) (Hernandez-Nicaise 1991; Takashima et al. 2013), consistent with a possible derivation from ancestral collar cells (Arendt et al. 2015).

Finally, the homology of the collar complex in animals and choanoflagellates is supported by its phylogenetic distribution. Its absence from all other eukaryotic lineages^2^ suggests that it is unlikely to evolve easily through convergence. Although there are subtle differences in the form and function of the collar of choanoflagellates and sponge choanocytes (Mah et al. 2014) (unsurprisingly, given that the two lineages diverged more than 600 million years ago), these differences are likelier to be the result of derivation from an ancestral collar cell than the similarities arising through convergent evolution. Therefore, bacterivorous collar cells likely trace their ancestry back to the choanozoan stem lineage, and the study of modern choanoflagellates holds the promise of illuminating the cellular foundations of animal origins.

## II. Division or aggregation? The two paths to multicellularity

How and when did the ancestors of animals become multicellular? Each of the multiple independent evolutionary origins of multicellularity (Figure 3) took place through one of two mutually exclusive mechanisms: clonal development, in which multicellularity arises through serial cell division without separation of sister cells, and aggregation, in which separate cells converge and adhere to each other. Clonal and aggregative multicellularity both have broad, scattered distributions in the phylogenetic tree of eukaryotes, suggesting that they both evolved a number of times independently (Figure 3A). Interestingly, the two pathways to multicellularity result in different types of multicellular forms, and arguably evolved in response to different selective pressures.

**Figure 3.**
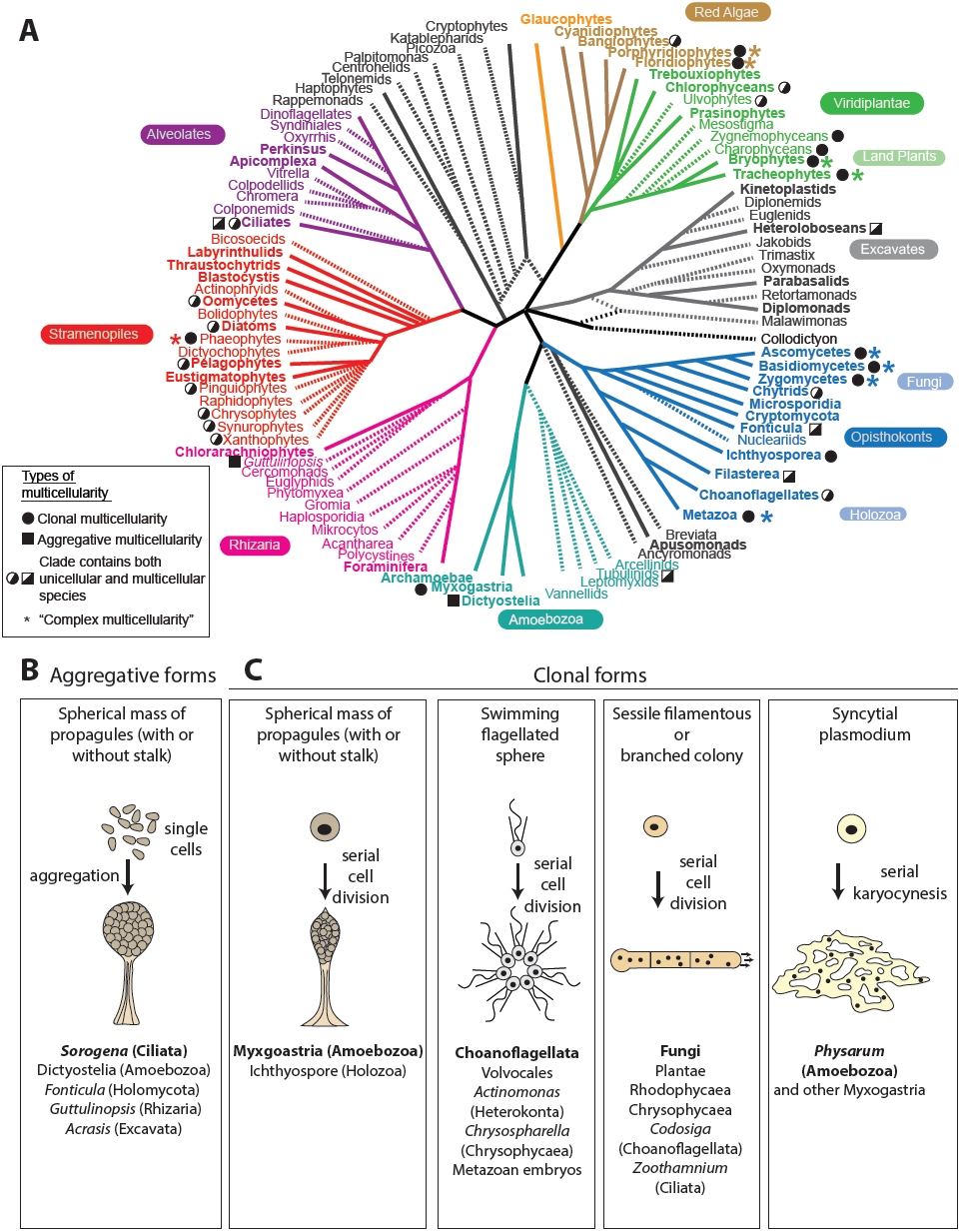
Clonal and aggregative multicellularity. (A) Phylogenetic distribution of clonal and aggregative multicellularity. Eukaryotic phylogeny is modified from (Keeling et al. 2014). Instances of multicellularity are mapped following (Bonner 1998; King 2004; Raven 2005; Ott et al. 2015; Sebé-Pedrós et al. 2017). (B and C) Examples of aggregative and clonal multicellularity. The organism indicated is shown in bold, and other organisms with similar forms of multicellularity are listed below. (B) Aggregative multicellularity gives rise to spherical masses of spores or cysts, sometimes atop a stalk. From (Bonner 1998; Brown et al. 2012; Du et al. 2015). (C) Clonal multicellularity gives rise to diverse multicellular forms. Drawn from (Bonner 1998; Fairclough et al. 2010).

### II.1. Independent origins of aggregative multicellularity

Although all extant animals display clonal development, could early animal ancestors have developed by aggregation? Though this hypothesis is hard to formally rule out, it appears unlikely in the current state of knowledge. The few cases of aggregation-like processes reported in animals are responses to artificial perturbations: for example, experimentally dissociated sponge cells can re-aggregate *in vitro* (Curtis 1962) but sponge embryonic development is strictly clonal *in vivo* (Ereskovsky 2010). We argue below that aggregative multicellularity is rarer than clonal multicellularity, and represents a distinct adaptive niche with a more limited evolutionary potential.

Aggregative multicellularity evolved at least seven times independently in eukaryotes (Figure 3A) as well as in some bacterial lineages such as Myxobacteria (Bonner 1998; Brown et al. 2012; Sebé-Pedrós et al. 2013b; Du et al. 2015). In nearly all well-characterized cases of aggregative multicellularity, cells respond to adverse conditions (e.g. starvation (Dworkin 1963; Stratford 1992; Souza et al. 1999)) by migrating toward each other and aggregating into a resistant mass of propagules (spores or cysts) – called a sorocarp, sporangium or (if a stalk is present) a fruiting body (Figure 3B) ^3^. The resulting structure is formed of quiescent cells that neither feed nor divide. Motility, if present (as in *Dictyostelium* slugs), is a transient step toward the formation of the sorocarp. The multicellular form eventually dissociates and the propagules disperse. Aggregation is thus a specialized “emergency response” in which cells transiently assemble under adverse conditions that compromise cell division, thus preventing clonal multicellularity. The potential advantages of aggregation include enhanced resistance to environmental stressors (with the outer cells shielding the inner cells from harmful agents such as UV and toxic chemicals (Stratford 1992)) and, in terrestrial species, formation of a stalk allowing elevation from the substrate and wind dispersal of the propagules (Bonner 1998).

Aggregation presents an impediment to the evolution of division of labor between cell types (Buss 1988): the multicellular mass is composed of cells that are not necessarily genetically related, making it especially vulnerable to invasion by “cheater” mutants that benefit from their presence in the aggregate without sharing resources or labor with the other cells (Strassmann et al. 2000; Santorelli et al. 2008). Indeed, theoretical arguments and experimental data suggest that aggregation can only be evolutionarily stable if it is restricted to closely related individuals (Gilbert et al. 2007; Kuzdzal-Fick et al. 2011). For example, the aggregative multicellularity of dictyostelids has been evolutionarily stable for more than 400 million years (Sucgang et al. 2011), and sophisticated ways to recognize kin have accordingly evolved (Benabentos et al. 2009; Hirose et al. 2011). Perhaps due to the difficulty of overcoming such conflicts, aggregative forms have a transient existence and little or no division of labor, with at most two to three coexisting cell types – usually germ cells (the inner spores or cysts) and somatic cells (e.g. the cells of stalk in the *Dictyostelium* fruiting body (Town et al. 1976; Dickinson et al. 2011)).

In all its forms, aggregative multicellularity involves an elaborate series of coordinated events: mutual cell attraction by chemotaxis, migration, adhesion, and, finally, differentiation into propagules (Du et al. 2015). How this cascade originated in each lineage is unknown in detail, but a likely evolutionary prerequisite was the presence of an inducible pathway for the sporulation/encystment of individual cells. Supporting this notion, comparison of dictyostelids and single-celled amoebozoans have shown that the cAMP/PKA pathway controlling the switch to multicellularity in dicytostelids evolved from an ancient encystment stress response found in diverse single-celled amoebae (Ritchie et al. 2008; Du et al. 2014; Kawabe et al. 2015). As it is contingent on specific prior, permissive evolutionary steps and vulnerable to cheaters, aggregative multicellularity might be both more difficult to evolve and less conducive to the evolution of differentiated cell types than is clonal multicellularity.

### II.2. Independent origins of clonal multicellularity

As clonal multicellularity can in principle result from a simple failure to complete cell division, it might evolve relatively easily through loss-of-function mutations (Bonner 1998). All three documented examples of multicellular forms that have evolved *de novo* in the laboratory developed clonally by incomplete cytokinesis within a small number of generations under selection (100 to 315; (Boraas et al. 1998; Ratcliff et al. 2012, 2013). The relative ease of evolving clonal multicellularity could explain why clonal forms are both phylogenetically more widespread (Figure 3A) and morphologically more diverse than aggregative forms. Indeed, clonal development underlies all known multicellular forms with active metabolism and proliferation (but also some sorocarps or fruiting bodies) (Figure 3C). Moreover, clonal multicellularity is likely less vulnerable to cheating than aggregation, as all cells share an identical genome (Buss 1988). These two factors together may account for the fact that all five known independently evolved instances of “complex multicellularity” (defined as obligate multicellularity with different cell types and elaborate organismal morphology (Knoll 2011)) involve clonal development: animals, fungi, green algae/land plants, red algae, and brown algae. The selective advantages that favored the evolution and maintenance of clonal multicellularity are unknown, but may include resistance to predators (Boraas et al. 1998), cooperative feeding (Koschwanez et al. 2011; Roper et al. 2013), simple division of labor (e.g. between flagellated cells and dividing cells) (Margulis 1981; Buss 1988; Michod 2007) and formation of a *milieu intérieur* (i.e. a controlled “internal environment” of stable composition). Finally, one cannot rule out the possibility that the initial evolution of animal multicellularity was a neutral event, that would have conferred no immediate fitness benefit and have first reached fixation by genetic drift.

## III. The origin of animal multicellularity

### III.1. How ancient is animal multicellularity?

The last common ancestor of animals was clearly multicellular, but might multicellularity extend back to the last common ancestor of choanozoans? Tantalizingly, many choanoflagellates form facultative multicellular forms, including swimming spherical (rosette), linear or flat colonies, and sessile branching colonies (Leadbeater 2014). These multicellular forms are all thought to develop clonally (Hibberd 1975; Karpov and Coupe 1998; Fairclough et al. 2010; Dayel et al. 2011; Leadbeater 2014). Although the phylogenetic distribution of known instances of multicellularity in choanoflagellates remains patchy, the life cycles of most species are incompletely known, precluding robust parsimony-based inference of the ancestral state. In favor of an ancient origin, the rosette colonies of three distantly related species (*Salpingoeca rosetta* (Leadbeater 1983; Fairclough et al. 2013), *Codosiga botrytis* (Hibberd 1975) and *Desmarella* Kent (Karpov and Coupe 1998)) are composed of cells joined by cytoplasmic bridges with a common distinctive ultrastructure featuring two parallel electron-dense plates flanking an electron-dense cytoplasmic core. The last common ancestor of these three species was also the last common ancestor of one of the two main choanoflagellate clades (Carr et al. 2008, 2017), suggesting that clonal multicellularity might already have evolved before the first split in the choanoflagellate phylogenetic tree. Characterization of the life cycles and multicellular development of more choanoflagellate species will further test this hypothesis. Whether clonal multicellularity arose once or several times in choanozoans, choanoflagellate rosettes offer a tantalizing proxy for the first stages of animal evolution, due to their close phylogenetic affinity to animals and to similarities in both cell ultrastructure and developmental mode.

Multicellularity might thus be as ancient as stem choanozoans, but could it be even older? The two closest relatives of choanozoans are filastereans and ichthyosporeans (Figure 1). Intriguingly, both form facultative multicellular forms, but in different ways from choanozoans – aggregation in filastereans (Sebé-Pedrós et al. 2013b) and fragmentation of a syncytium in ichthyosporeans (Suga and Ruiz-Trillo 2013). If holozoan multicellular forms are homologous to each other, this would imply that interconversions took place between aggregative and clonal multicellularity in stem-filastereans or stem-choanozoans^4^. Perhaps more likely is the possibility that multicellularity evolved independently in the three holozoan clades, although this remains to be tested.

### III.2. The genetic basis for the origin of animal multicellularity

What molecular mechanisms first supported the evolution of animal multicellularity? This problem cannot be studied solely in animals, as there is no known animal mutant where cleavage of the zygote would directly produce separate free-living cells rather than an embryo, thus reverting to unicellularity. To answer this question, it is therefore necessary to study phylogenetically relevant groups with facultative multicellularity. Thus far, a dozen genes have been found to be either necessary or sufficient for multicellularity in the four groups investigated: green algae, fungi, slime molds and choanoflagellates^5^. All known multicellularity genes encode proteins that belong to one of two major functional categories: extracellular matrix (ECM) proteins and, in the case of clonal multicellularity, cytokinesis regulators (Table 1). This suggests that the initial evolution of multicellularity on different branches of the tree of life repeatedly converged on similar mechanisms (Abedin and King 2010).

Among the taxa studied, choanoflagellates occupy a privileged position as the sister-group of animals. The recent establishment of forward genetics in the model choanoflagellate *Salpingoeca rosetta* has revealed the first gene known to be required for multicellular development, named *rosetteless* for its mutant phenotype (Levin et al. 2014). While wild-type *S. rosetta* reliably develops into spherical colonies (called “rosettes”) upon induction with bacterial signals (Dayel et al. 2011; Alegado et al. 2012; Woznica et al. 2016), *rosetteless* mutants are unable to form rosettes under all studied conditions (although the ability to develop into another clonal multicellular form, linear chains, is unaffected). The *rosetteless* gene encodes a C-type lectin that is secreted into the core of the rosette as an ECM component (Figure 4A). In *S. rosetta*, the integrity of the colonies is thus possibly ensured by the basal extracellular matrix, to which cells appear to anchor by filopodia (Dayel et al. 2011). Both ECM and filopodia also contribute to the cohesion of animal blastulae. Blastomeres are held together by an abundant ECM rich in lectins (Fraser and Zalik 1977; Roberson and Barondes 1983; Harris and Zalik 1985; Outenreath et al. 1988; Lee et al. 1997) and appear linked by filopodia in sponges (Ereskovsky 2010), cnidarians (Benayahu et al. 1989), echinoderms (Vacquier 1968), amphioxus (Hirakow and Kajita 1994), and mouse (Salas-Vidal and Lomeli 2004). In mice, laser ablation of the filopodia results in loss of blastomere cohesion (Fierro-González et al. 2013). Thus, the multicellular state of choanoflagellate rosettes and animal embryos is established and maintained by comparable mechanisms.

**Figure 4.**
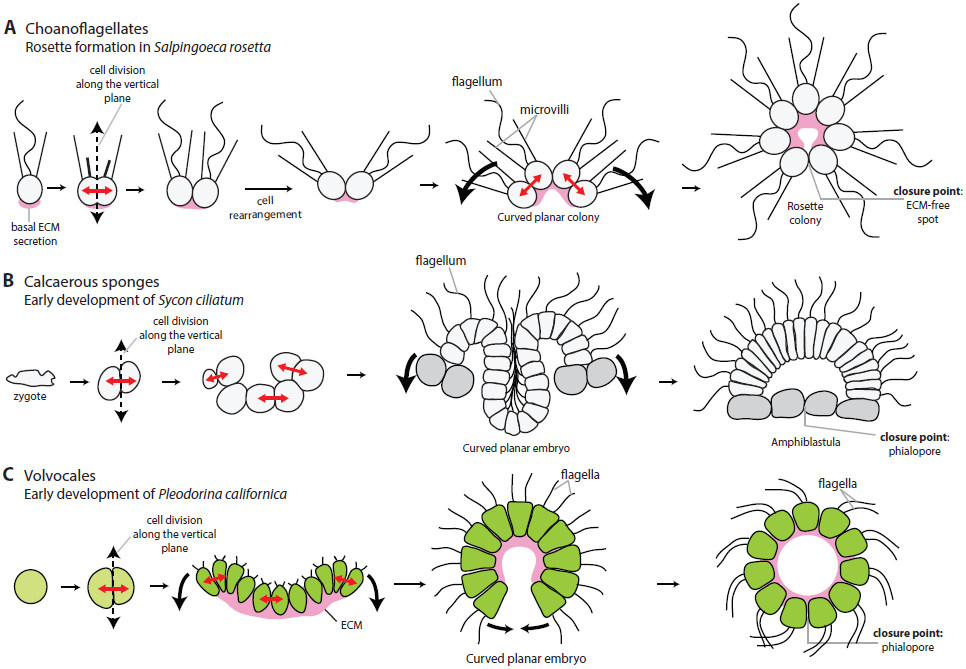
Morphogenesis in choanoflagellate rosettes, calcareous sponge embryos and volvocale embryos. (A) Morphogenesis during rosette formation in the choanoflagellate *Salpingoeca rosetta*, following (Fairclough et al. 2010). (B) Early embryonic development of the calcareous sponge *Sycon ciliatum*, including amphiblastula inversion, from (Franzen 1988). (C) Early embryonic development of the volvocale *Pleodorina californica* (Höhn and Hallmann 2016). In other volvocales such as *Volvox*, an additional developmental stage is intercalated, in which the embryo first forms a sphere with flagella pointing inward, which later opens up into a sheet before inversion.

An intriguing open question is whether cadherins, which contribute to adhesion between animal embryonic and adult cells (Abedin and King 2010), are also involved in rosette formation. Choanoflagellate genomes encode a large number of cadherins (23 in *Monosiga brevicollis* and 29 in *S. rosetta*) (Nichols et al. 2012) but their localization is so far only known in the solitary species *Monosiga brevicollis*, in which the cadherins MBCDH1 and MBCDH2 were found localized to the collar and might contribute either to collar integrity or to bacterial capture (Abedin and King 2008).

### III.3. Choanoflagellate multicellularity and the origin of animal embryogenesis

The similarity of choanoflagellate rosettes to the blastula/morula stage of animal development is consistent with a modern version of Haeckel’s Blastaea hypothesis – which proposes that early animal ancestors were motile spheres of flagellated cells that formed clonally (Haeckel 1874, 1892; Nielsen 2008; Arendt et al. 2015). Indeed, the first developmental stage of marine invertebrates is often a ciliated free-swimming blastula that resembles choanoflagellate rosettes (minus the microvillar collar) and, given its widespread taxonomic distribution, was plausibly part of development in the Urmetazoan (Nielsen 2012).

How did serial cell divisions give rise to these early multicellular forms? In most eukaryotes, including choanoflagellates, cell division has to accommodate a constraint: the two centrioles that form the flagellar basal body during interphase also organize the mitotic spindle during division. Moreover, in choanoflagellates, microvilli are directly transmitted to daughter cells. As a consequence, the plane of cell division in choanoflagellates must comprise the apical pole, where centrioles and microvilli are located, and symmetric cell divisions necessarily take place along the apico-basal axis (Figure 4A). This constraint on the direction of cell division seems universal in choanoflagellates (Leadbeater 2014; Anderson et al. 2016). As a consequence of this fixed division orientation, in the absence of cell reorientation and/or rearrangements, cell proliferation alone can only produce cells linked together in linear chains (e.g. in *S. rosetta* (Dayel et al. 2011)) or planar sheets (e.g. in *Choanoeca perplexa* – formerly *Proterospongia choanojuncta* (Leadbeater 1983)). Any more complex shape – such as a spherical rosette – must be achieved by cell rearrangements after division to allow bending and, ultimately, “closure” of the sheet at the point where non-sister cells meet.

These cell rearrangements are apparent during rosette formation in *S. rosetta* (Fairclough et al. 2010) and the final “closure point” can be identified in mature rosettes as an ECM-free spot visible in immunostainings for the Rosetteless protein (Woznica et al. 2016). This constraint seems to apply to any spherical colony formed by clonal division of motile flagellated cells: in flagellated volvocale green algae, a similar folding process takes place (Höhn and Hallmann 2016). It plausibly applied to the first animal embryos as well: indeed, a similar series of steps takes place during the early development of calcareous sponges, which first form as concave sheets and secondarily fold into spherical embryos called “amphiblastulae” (Figure 4B) (Franzen 1988; Ereskovsky 2010; Arendt et al. 2015). In both volvocales and calcareans, the hole resorbed at the closure point is called the phialopore. In other animals (eumetazoans and non-calcareous sponges), after the evolution of non-flagellated zygotes, this constraint was eventually overcome: early cell division can take place along all cellular axes and cleavage directly produces a spherical blastula (Arendt and Nübler-Jung 1997).

## IV. The ancestry of animal cell differentiation

### IV.1. Alternative hypotheses for the origin of animal cell types: temporal-to-spatial transition and division of labor

Choanoflagellate rosettes and the multicellular forms of ichthyosporeans and filastereans are not known to undergo cell differentiation. This contrasts with the “complex multicellularity” (Knoll 2011) of animals, which are mosaics of different cell types with controlled spatial distribution. The origin of cell types thus represents one of the key steps in the evolution of animals from their microeukaryote ancestors. Two main hypotheses have been put forward to explain the evolution of animal cell differentiation: the temporal-to-spatial transition (TST) hypothesis (Zakhvatkin 1949; Mikhailov et al. 2009; Sebé-Pedrós et al. 2017) and the division of labor (DOL) hypothesis (Mackie 1970; Arendt 2008). The TST hypothesis proposes that cell differentiation predated multicellularity, building upon the observation that many modern microbial eukaryotes can switch between different cell phenotypes over their life history (Figure 5). In this scenario, some of these temporally alternating phenotypes were converted into spatially segregated cell types in animal ancestors. The DOL hypothesis starts from the observation that individual microbial eukaryotes combine in a single cell multiple functions that are accomplished by different cell types in animals (Figure 6), including the perception and transduction of environmental signals, movement, feeding, and division. In the DOL hypothesis, the bulk of cell differentiation evolved after multicellularity, by differential loss of function from multifunctional ancestral cell types. The TST and the DOL hypotheses are complementary rather than mutually exclusive, and it is plausible that both processes contributed to the early evolution of animal cell types.

**Figure 5.**
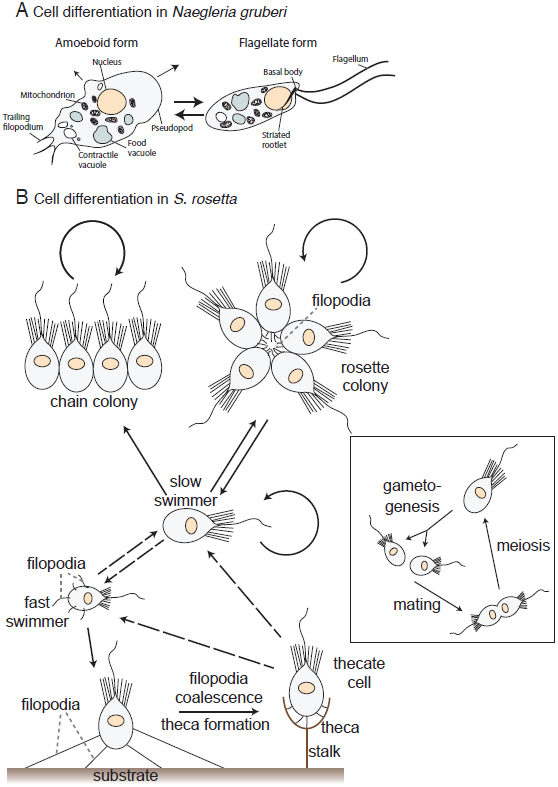
Temporally alternating cell types in protozoans. (A) The heterolobosean excavate *Naegleria gruberi* can switch between a flagellated swimmer phenotype and a deformable crawler (“amoeboid”) phenotype. Redrawn from (Fritz-Laylin et al. 2010). (B) Cell types and life history transitions in the choanoflagellate *S. rosetta*, from (Dayel et al. 2011; Levin and King 2013). Main panel depicts the dynamic asexual life history of *S. rosetta* whereas the inset indicates its sexual cycle. Dotted lines indicated inferred transitions that have not been directly observed.

**Figure 6.**
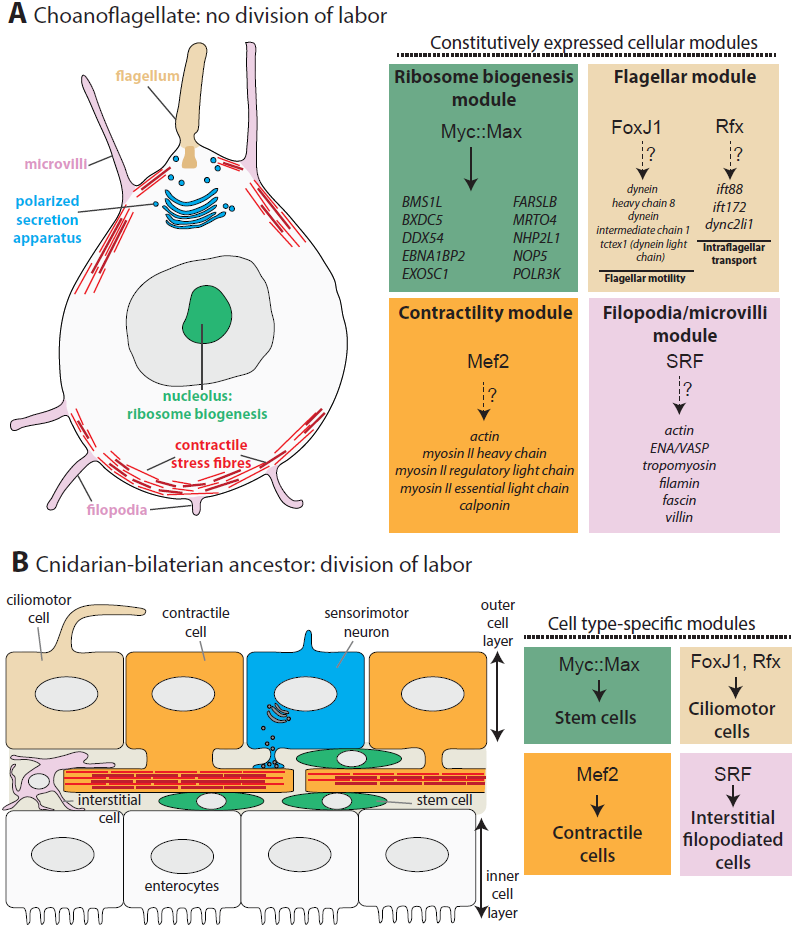
The division of labor hypothesis. (A) Cellular modules present in the choanoflagellate *Salpingoeca rosetta*. On the right: choanoflagellate orthologs of the selector transcription factors that control these modules in animals, on top of a list of choanoflagellate orthologs of their animal targets (Supplementary Figure 3 and references in the text). The transcription factors are FoxJ1 and RFX for flagella (Stubbs et al. 2008; Yu et al. 2008; Thomas et al. 2010; Vincensini et al. 2011; Vij et al. 2012), Mef2 for contractility (Black and Olson 1998; Sebé-Pedrós et al. 2011; Brunet et al. 2016), SRF for filopodia (Knöll and Nordheim 2009; Gervais and Casanova 2011; Weinl et al. 2013; Franco et al. 2013), and the Myc::Max complex for stem cells (Young et al. 2011). No terminal selector is indicated for the secretion apparatus, as there seems to be no known neural terminal selector with a choanoflagellate ortholog. Dotted lines indicate that it is unknown whether the choanoflagellate transcription factors control the same genes as their animal orthologs, except for the Myc::Max complex for which regulation is indicated by computational predictions (Brown et al. 2008) and electrophoretic mobility shift assays (Young et al. 2011). (B) Cellular modules shown in A are segregated into distinct cell types in animals (here, the putative cell type complement of stem-eumetazoans is illustrated based on the cell types shared by cnidarians and bilaterians (Fautin and Mariscal 1991; Schmidt-Rhaesa 2007; Arendt et al. 2015)) and terminal selector transcription factors specify distinct cell types.

### IV.2. Testing the TST hypothesis

Many single-celled eukaryotes have temporally alternating cell phenotypes (Figure 5). The distinct morphologies, transcriptomes (Fairclough et al. 2013; Sebé-Pedrós et al. 2013b; de Mendoza et al. 2015), proteomes (Sebé-Pedrós et al. 2016b), and chromatin states (Sebé-Pedrós et al. 2016a) of these cell phenotypes in single-celled holozoans suggest that they represent stable cell types, like those existing in animals, rather than instances of short-term phenotypic plasticity. Are some of these cell types homologous to those of animals, as the TST hypothesis postulates? The simplest example of temporal-to-spatial transition is probably meiosis, which clearly predated animals and dates back to the last common eukaryotic ancestor (Ramesh et al. 2005). In choanoflagellates, gametes competent to undergo cell-cell fusion directly transdifferentiate from haploid solitary cells (Figure 5B) (Levin and King 2013; Woznica et al. 2017), while in all extant animals meiosis and gametogenesis start from diploid cells integrated within the adult organism (Nielsen 2012).

Another temporal switch frequently found in protozoa that has been proposed to have been co-opted in animal cell differentiation is the alternation between a flagellated phenotype (allowing locomotion in an aqueous environment by flagellar beating) and a deformable “amoeboid” crawling phenotype that navigates solid environments by actin-mediated deformations of the cell body^6^ (Figure 5A). These two phenotypes and associated genes are broadly distributed in the eukaryotic tree of life (Supplementary Figure 2), suggesting that the last common eukaryotic ancestor might have been able to switch between both (Fritz-Laylin et al. 2010, 2017). If so, this switch could have existed in the last single-celled ancestors of animals^7^. Suggestively, virtually all animals combine static epithelial cells (often bearing a flagellum and lining the body surface) with deformable crawling cells located in the primary body cavity that patrol the tissues by actin-mediated locomotion. These migratory interstitial cells usually function as phagocytes (and sometimes also as stem cells) and quickly accumulate upon infection, wound repair or allograft rejection. They are called archaeocytes in sponges (Cheng et al. 1968; Alié et al. 2015), amoebocytes in cnidarians (Patterson and Landolt 1979; Olano and Bigger 2000; Couch et al. 2013) and hemocytes or macrophages in bilaterians (Hartenstein 2006; Schmidt-Rhaesa 2007). If the interstitial phagocytes of these large animal clades are homologous (which remains to be tested), they could have evolved from the crawling phase of the Urchoanozoan or one of its descendants (Mendoza et al. 2002; Arendt et al. 2015). In parallel, the flagellated phase of this organism would have provided the genetic program for the ciliated epithelium that putatively covered the surface of the Blastaea and subsequently diversified into several epithelial cell types and into other cell types thought to have evolved from epithelia (epidermal cells, enterocytes, sensory cells, neurons and muscle cells (see section IV.3 and Figure 6B)).

How could one test the homology between animal and single-celled holozoan cell types? In the past few years, the comparison of transcriptomes has emerged as a promising approach to investigate cell type homology (Arendt 2005, 2008; Lauri et al. 2014) and to build cell type trees (Liang et al. 2015; Kin et al. 2015; Musser and Wagner 2015). Central to these comparisons are the “terminal selector” transcription factors that directly implement cell phenotypes by sitting directly above large batteries of differentiation genes (Arendt et al. 2016; Hobert 2016). This approach can potentially be extended to single-celled holozoans: do terminal selector transcription factors play a role in establishing and maintaining their cell types? If so, how do they compare to their animal counterparts? These questions are still open, but intriguing potential case studies can already be identified. For deformable crawlers, an interesting candidate is the Runx family of transcription factors (Coffman 2003), which were likely present as a single copy in urmetazoan and urholozoan progenitors (Rennert et al. 2003; Sullivan et al. 2008; de Mendoza et al. 2013). Runx transcription factors specify circulating cells that move by actin-mediated crawling (Pancer et al. 1999; Otto et al. 2003; Waltzer et al. 2003; Burns et al. 2005) and directly promote cell motility (Leong et al. 2010; Zusso et al. 2012; Lie-A-Ling et al. 2014; VanOudenhove et al. 2016). High levels of Runx transcription have been detected by RNA-seq in the archaeocytes of the sponge *Ephydatia fluviatilis* (Alié et al. 2015), suggesting that the link between this transcription factor family and the crawling cell phenotype might be ancient in animals. Runx is also present in the genomes of ichthyosporeans and filastereans (de Mendoza et al. 2013), which both have a crawling (“amoeboid”) phase. The *runx* gene is significantly upregulated during the crawling phase of the ichthyosporean *Creolimax* (de Mendoza et al. 2015) and, in *Capsaspora*, computationally predicted Runx targets show a significant enrichment for genes encoding actin cytoskeleton and other proteins involved in crawling motility (Sebé-Pedrós et al. 2016a).

Another potentially relevant transcription factor with stem-holozoan ancestry is Brachyury, which is upregulated in *Capsaspora* amoebae and is predicted to regulate homologs of genes controlled by Brachyury in mouse, including those required for cell motility (Sebé-Pedrós et al. 2016a). In animals, Brachyury is often involved in the motility of embryonic (but not adult) cells (Yanagisawa et al. 1981; Gross and McClay 2001). Both Runx and Brachyury appear to have been lost from sequenced choanoflagellate genomes, in line with the possibility that choanoflagellates might have lost crawling by whole-cell deformation. Direct mechanistic studies of these and other transcription factors in both single-celled holozoans and non-bilaterian animals, as well as more extensive molecular characterization of the relevant cell types in a broad sampling of phylogenetically relevant species, will help in testing these hypotheses and in revealing other potential instances of TST.

### IV.3. Testing the DOL hypothesis

The phenotypic comparison of animal cells with free-living microeukaryotes offers some direct observations in support of the DOL hypothesis. Several cellular features that are constitutively present in choanoflagellates are restricted to a subset of animal cell types. These include a motile flagellum, filopodia, contractile stress fibers, polarized secretion (Burkhardt et al. 2011), and mitotic machinery – respectively restricted in animals to ciliated cells (in humans: vestibular hair cells, spermatozoa, tracheal lining, fallopian tubes), migratory cells, muscle cells (and other contractile cells), neurons (and other secretory cells), and stem/progenitor cells. In support of the antiquity of these cellular modules, many of the effector genes that implement these phenotypes in animals are conserved in choanoflagellates and other holozoans (Figure 6A). Intriguingly, the same is true of the selector transcription factors that control expression of these modules in animals (Figure 6A) – for example FoxJ1 and RFX for flagella (see Figure 6 for other examples and the legend for references). It is currently unknown whether the targets of most of these transcription factors are the same in choanoflagellates as in animals. In the case of Myc, bioinformatic data suggest that the choanoflagellate ortholog preferentially controls ribosomal biogenesis genes, as it does in animals (in line with the general requirement for intense protein synthesis during proliferation) (Brown et al. 2008), and that RFX controls at least a few flagellar genes in *Monosiga brevicollis* (Piasecki et al. 2010).

Detailed comparisons of these cellular modules and of their upstream transcriptional circuits with the cell type-specific modules and circuits of animals will be crucial to further testing the DOL hypothesis. The targets of the cell type-specific transcription factors (Figure 6) could be determined experimentally by ChIP-seq and/or loss-of-function experiments in choanoflagellates and other single-celled holozoans. If they are at the top of the same regulatory networks as in animals, this would further support the DOL hypothesis, and would suggest a mechanistic basis for division of labor. Indeed, it would imply that the transcriptional networks of the single-celled ancestors of choanozoans or holozoans were modular, with different transcription factors sitting upstream of functionally distinct gene modules. Such an architecture would imply that animal ancestors were ‘pre-adapted’ to evolving division of labor once multicellularity evolved, as a module could be selectively ‘shut down’ in a given cell by inhibiting its upstream transcription factor with minimal impact on the expression of other modules. Intriguingly, and consistent with this hypothesis, a modular architecture of gene regulatory networks, with functionally related genes or operons being controlled by the same transcription factors, has repeatedly been found in single-celled organisms including *E. coli*, *B. subtilis* (Shen-Orr et al. 2002; Madan Babu and Teichmann 2003; Fadda et al. 2009) and yeast (Tavazoie et al. 1999; Tanay et al. 2004). The advantage of such a modular architecture in single-celled organisms is not generally known, but possibilities include modulating the total synthesized quantity of a module (depending for example on environmental signals or on the phase of the cell cycle), ensuring stoichiometry between its components, or allowing regeneration of a given part of the cell. For example, choanoflagellates regenerate their flagellum after each division and can also do so when recovering from microtubule-depolymerizing treatment (Froesler and Leadbeater 2009). In the unicellular alga *Chlamydomonas*, flagellum regeneration involves a coordinated rise in flagellar gene transcription (Keller et al. 1984) and investigation of promoter motifs suggests shared regulation of these flagellar genes by yet unidentified, specific transcription factors (Stolc et al. 2005).

Currently, the evidence for the TST and DOL scenarios remains primarily descriptive and restricted to a few candidate genes. It will be crucial to ground future comparative studies in a firm mechanistic basis to more fully test these two hypotheses and investigate their respective contributions to the origin of animal cell types. This endeavor should benefit from ongoing efforts to obtain an unbiased transcriptomic, proteomic and phenotypic characterization of animal cell types, notably by single-cell approaches (Achim et al. 2015; Wagner et al. 2016; Villani et al. 2017; Regev et al. 2017). Analyzed with refined methods of phylogenetic reconstruction and ancestral state inference (Liang et al. 2015; Kin et al. 2015), such datasets should help illuminate our understanding of the origin and evolution of animal cell differentiation.

## Conclusion and outlooks

In the pre-molecular era, the reliance on morphology to reconstruct the origin of animals posed a risk of circular reasoning, as ancestral state reconstructions relied on phylogenetic trees that were themselves built from hypotheses of morphological evolution. This circularity was broken with the application of molecular phylogenetics, which has resolved the relationships among most animal phyla to each other and to protozoan relatives, with only a few outstanding controversies – such as whether the sponge or ctenophore lineage branched first (Whelan et al. 2015; Simion et al. 2017). By mapping morphological features onto molecular phylogenies, it has become possible to re-approach the morphology, cell biology and development of the first animals from a modern perspective.

The first animals likely developed through serial division of flagellated, bacterivorous cells that sported a microvillar collar. Diverse cell types then evolved through a combination of innovation, division of labor, and spatiotemporal regulation of cell types that were ancestrally separated temporally in life histories of the progenitors of animals. Determining the relative contributions of these different mechanisms, and the interrelationships among different cell types in diverse animals will not only help us understand our evolutionary origins, but help to deepen our understanding of the structure and function of animal cells themselves. Indeed, insights into evolutionary relationships have already led to unexpected discoveries in cell biology – e.g., the discovery that parasitic apicomplexans such as *Toxoplasma* and *Plasmodium* harbored modified plastids provided the rationale for targeting them with drugs known to affect cyanobacteria (Fichera and Roos 1997); and comparative characterizations of ciliary proteomes have, by identifying an evolutionarily conserved core, revealed central players of clinical relevance (Li et al. 2004).

In the future, it will be valuable to develop functional molecular and cellular techniques in a broader range of early-branching animals and single-celled holozoans, and to expand observational research to clarify uncertainties concerning, for example, the life cycles of additional choanoflagellates or the embryology of sponges and placozoans. Transcriptome sequencing of diverse cell types from choanoflagellates and early branching animals may also help to reveal the relative importance of division of labor as opposed to temporal-to-spatial transitions in the evolution of animal cell differentiation. By integrating these lines of research, we hope to gain a clearer picture of how our protozoan ancestors emancipated from the microbial world and founded the animal kingdom.

## Acknowledgements

We thank all members of the King lab for discussion of the ideas presented, David Booth, Ben Larson and Monika Sigg for critical reading of the manuscript, Patrick Keeling for sharing the figure panel on eukaryote phylogeny, and Daniel J. Richter and Cédric Berney for discussions on defining Choanozoa.

---

**Supplementary Figure 1. Collar cells of cnidarians and bilaterians.** All panels are TEM. (A) Epidermal collar cell from the ciliary bands of the brachiolaria larva of the starfish *Asterina rubens*, from (Nerrevang and Wingstrand 1970). (B) Protonephridial terminal cell from the trochophore larva of the annelid *Glycera* (Ruppert and Smith 1988). (C) Gastrodermal collar cell from the cnidarian *Polypodium hydriforme* (Raikova 1995). (fl): flagellum, (mv): microvilli, (bb): basal body, (vc): vacuole, (mu): mucus.

**Supplementary Figure 2. Phylogenetic distribution of flagellated swimmers and of deformable crawling (“amoeboid”) cells.** Eukaryotic phylogeny is modified from (Keeling et al. 2014). Distribution of crawling cells follows (Fritz-Laylin et al. 2017). Cells with pseudopodia used for prey capture and phagocytosis but not for motility are not classified as deformable crawlers (e.g. choanoflagellates (Leadbeater 2014), parabasalids (Brugerolle 2005)).

**Supplementary Figure 3. SRF and FoxJ phylogenetic trees.** (A) FoxJ family phylogenetic tree. The split between the FoxJ1 clade and the FoxJ2/3 clades is inferred to have preceded choanozoans, in accordance with previous reports (Larroux et al. 2008). Two Forkhead-box proteins which are mutual best BLAST hits with *Homo sapiens* FoxJ1 cluster in a clade together with animal FoxJ1. (B) SRF family phylogenetic tree. The tree is rooted with the Mef2 family, the most closely related family of MADS-box transcription factors (including a previously identified ortholog in *Monosiga brevicollis* (Sebé-Pedrós et al. 2011) and a newly predicted ortholog in *Salpingoeca rosetta* on genomic scaffold supercontig 1.36 positions 82482-84459). A *Monosiga brevicollis* predicted protein annotated as SRF and the *Salpingoeca rosetta* protein PTSG_11095 (both mutual best BLAST hits with *Homo sapiens* SRF) cluster in a clade with animal SRF. Both trees have been reconstructed with MrBayes 3.2.3, with 10,000 generations under the GTR substitution model with default assumptions and convergence assessed by standard deviation <0.01. Support values are displayed on nodes.

## Footnotes

1 Although the collar complex appears choanozoan-specific, microvilli in general may be of more ancient origin. The protozoan *Ministeria vibrans*, which belongs to Filasterea (the sister-group of choanozoans), displays microvilli-like radiating tentacles that are distributed over the whole cell cortex and are of constant length within a given cell at a given time (Cavalier-Smith and Chao 2006). Neither the ultrastructure of the radiating tentacles nor presence of ERM and Whirlin homologs in the *Ministeria* genome have yet been reported.

2 Although morphologically analogous forms exist in a few distantly related species (Mah et al. 2014), in all cases the underlying ultrastructure differs. For example, in the amoebozoan *Phalansterium* (Cavalier-Smith et al. 2004), the flagellum is surrounded by a “collar” formed by a continuous fold of cytoplasm rather than by independent microvilli (Hibberd 1983). In the pedinellales (a group of heterokonts (Cavalier-Smith and Chao 2006)) *Actinomonas* and *Pteridomonas*, the flagellum is surrounded by a double ring of tentacles which are supported by microtubule triads rather than actin microfilaments, and are thus not microvilli (Larsen 1985; Patterson and Fenchel 1985). Finally, the golden alga *Chrysosphaerella multispina* (Bradley 1964) has a superficially collar cell-like appearance, with a flagellum flanked by two spines of acellular silicate (Kristiansen 1969) instead of microvilli.

3 The few exceptions represent an infrequently observed type of aggregative multicellularity, sexual agglutination, that has been reported in yeasts (Wickerham 1958), choanoflagellates (Woznica et al. 2017) and *Chlamydomonas* (Bergman et al. 1975). Like other cases of aggregation, it results in transient multicellular forms without coordinated motility, feeding or proliferation, and which dissociate after karyogamy has been accomplished. In some species (such as *Chlamydomonas*) environmental stress (e.g. starvation) is a precondition to sexual agglutination.

4 There is no known definitive example of such a transition. The most plausible candidates are slime molds, in which similar fruiting bodies form clonally in Myxogastria and by aggregation in Dicytostelia (Stephenson and Schnittler 2016) which might be sister-groups (Fiore-Donno et al. 2010). However, no individual species is known to be able to develop multicellularity alternatively by either route.

5 Some pleiotropic genes are necessary for multicellularity as an indirect effect of them being involved in a more general cell function, such as transcription or cell motility. For example, across the 123 *Dictyostelium* mutants with aberrant or abolished aggregation (Glöckner et al. 2016), most are deficient in transcription, cell movement or cAMP synthesis. These genes are not discussed here as their role is indirect.

6 In the past, shape-shifting and crawling cells have often been encompassed within the general adjective “amoeboid” (Taylor and Condeelis 1979). More recently, there has been a tendency to distinguish different types of deformable crawlers based on the mechanisms of movement – restricting the term “amoeboid migration” to fast-migrating cells moving by actomyosin contraction and blebbing independently of adhesion (Liu et al. 2015; Ruprecht et al. 2015). Other categories include “mesenchymal migration” for slow-migrating animal cells with a lamellipodium propelled by actin polymerization and integrin adhesion (Panková et al. 2010) and more recently “αmotility” for fast migration with a pseudopod propelled by actin polymerization without specific adhesion (Fritz-Laylin et al. 2017). A common feature of nearly all deformable crawlers is their reliance on actin microfilaments (the only known exception is nematode sperm (Bottino et al. 2002)). Here, in line with recent studies (Fritz-Laylin et al. 2017), we use the general term “crawling cells” to refer globally to these cells regardless of the precise underlying mechanism.

7 Notably, no deformable crawling phase is known in extant choanoflagellates (despite one early unconfirmed report (Saville-Kent 1880)), suggesting they might have lost it – or that it does not occur under standard laboratory conditions. Choanoflagellates do show the ability to crawl on solid substrates by dynamic extension and retraction of adhesive filopodia prior to settlement, but without apparent deformations of the cell body outside filopodia (Figure 5B) (Dayel et al. 2011).

